# Metagenomic evidence for co-occurrence of antibiotic, biocide and metal resistance genes in pigs

**DOI:** 10.1101/2021.01.26.428250

**Authors:** Xuanji Li, Christopher Rensing, Gisle Vestergaard, Joseph Nesme, Shashank Gupta, Manimozhiyan Arumugam, Asker Daniel Brejnrod, Søren Johannes Sørensen

**Author notes:** **Corresponding author**: Asker Daniel Brejnrod & Søren J. Sørensen, Universitetsparken 15, bldg. 1, DK2100 Copenhagen.

## Abstract

Antibiotic-resistant pathogens constitute an escalating public health concern. Hence a better understanding of the underlying processes responsible for this expansion is urgently needed. Co-selection of heavy metals/biocides and antibiotic resistance genes (ARGs) has been suggested as one potential mechanism promoting the proliferation of antimicrobial resistance (AMR). This paper aims to elucidate this interplay and exploit differences in antibiotic usage to infer patterns of co-selection by the non-antibiotic factors metals and biocides in the context of pig farming. We examined 278 gut metagenomes from pigs with continuous antibiotic exposure, only at weaning and at no exposure. Metals as growth promoters and biocides as disinfectants are currently used with little restrictions in stock farming. The pigs under continuous antibiotic exposure displayed the highest co-occurrence of ARGs and other genetic elements while the pigs under limited use of antibiotics still showed abundant co-occurrences. Pathogens belonging to *Enterobacteriaceae* displayed increased co-occurrence phenomena, suggesting that this maintenance is not a random selection process from a mobilized pool but pertains to specific phylogenetic clades. These results suggest that metals and biocides displayed strong selective pressures on ARGs exerted by intensive farming, regardless of the current use of antibiotics.

**Highlights:**

- A comprehensive gut microbiome metagenomics analysis of 278 pigs
- Co-selection phenomena were investigated via co-occurrence patterns as a proxy
- Twenty-seven types of co-occurrences involving 131 resistance genes were detected
- Regardless of use of antibiotics, AMR can be maintained by co-occurrence with MRGs/BRGs
- Maintenance of AMR is not a random selection process but pertains to specific phylogenetic clades

## Introduction

Antimicrobial resistance in pathogenic bacteria constitutes a considerable public health concern (Baker-Austin et al., 2006). Despite making tremendous efforts to restrict usage of several key antibiotics worldwide, AMR can frequently be found in different environments (B. Li et al., 2015; Rodriguez-Mozaz et al., 2015). To control the potential threat from AMR spread, it is necessary to investigate the underlying processes responsible for this expansion.

Generally, the occurrence of AMR is largely driven by the selection pressure of antibiotics (Goossens et al., 2005). Besides that, other agents such as antibacterial biocides and heavy metals can help to maintain AMR via co-selection (Ashbolt et al., 2013; Hartmann et al., 2016; Mazhar et al., 2021). The co-selection of ARGs and metal resistance genes (MRGs) has been commonly observed in contaminated soil (Song et al., 2017), water (Icgen and Yilmaz, 2014), and industrial environments (Mazhar et al., 2021; Stepanauskas et al., 2005) with enriched metals. Some literature has demonstrated that ARGs had a stronger correlation to some metals than to their corresponding antibiotics (Ji et al., 2012). Unlike antibiotics, metals can constitute a long-term selection pressure due to metals not being degradable (Stepanauskas et al., 2005), which can bring stability to the ARG pool in both engineered and natural systems. Compared to metals, biocides have received less attention as a potential co-selection agent. Biocide substances have been employed for centuries as disinfectants for medical equipment and as preservatives in pharmaceuticals, cosmetics and food (SCENIHR, 2009), especially biocides with limited toxicity to animals such as quaternary ammonium compounds (QACs), biguanides and bisphenols (Grande Burgos et al., 2013). However, some research has suggested excessive use of QACs increased bacterial tolerance to antibiotics (Buffet-Bataillon et al., 2012; Tandukar et al., 2013). Metals and biocides can co-select for antibiotic-resistant bacteria by several mechanisms (Chapman, 2003): resistance genes are physically located on the same genetic element (co-resistance); the same gene confers resistance to both antibiotics and biocide/metal (cross-resistance); biocide resistance genes (BRGs)/MRGs share the same regulatory mechanism with ARGs (co-regulation).

The concurrent transfer of ARGs and BRGs/MRGs via mobile genetic elements (MGEs) is another challenging public health problem, which has aggravated the persistence and spread of ARGs (Benveniste and Davies, 1973). So far, research on the co-transfer of these three genetic determinants has been limited, although there are plenty of opportunities for this to occur. Stock farming is one such potential hot spot for the dissemination of combinations of ARGs, BRGs/MRGs, and MGEs due to the widespread use of antibiotics, heavy metals and biocides in food feed (Clark, 2004). Antibiotics and metals (especially zinc and copper) are commonly used as growth promoters in stock farming (Gaskins et al., 2002). However, the use of antibiotics as growth promoter has been banned in the European Union since 2006 (Lekagul et al., 2019) and therefore farmers have given more attention to alternatives such as copper and zinc (Jensen et al., 2016). This has resulted in the accumulation of these metals in soils where pig slurry was applied. In contrast, China does not prohibit the use of antibiotics as growth promoter (Lekagul et al., 2019) in stock farming and at the same time, the extensive use of heavy metals in feed in China has greatly increased over the last couple of decades (Wang et al., 2013). Many of the plants used for livestock are dependent on biocides to control weeds, insects and diseases (Clark, 2004) and biocides are also used for disinfection of animal housing facilities (Montfoort JA, Poel P van der, n.d.). However, investigations on the co-selection of heavy metals/biocides and antibiotics used in animal farms, which are closely connected to human health, are still scarce (Seiler and Berendonk, 2012), especially under various exposure levels of antibiotics.

Genome sequencing has often complemented analysis of phenotypic outcomes (Song et al., 2017; Stepanauskas et al., 2006) to clarify the genetic determinants of resistance, and indeed large-scale efforts have investigated repositories of publicly available genomic data and established the co-presence of both metal- and antibiotic-resistance (Li et al., 2017; Pal et al., 2015). These repositories are comprehensive, but the information about the origins of the sequenced isolates varies, making it difficult to infer that genes of interest are present in the underlying populations as a whole. In this study, we attempted to make use of publicly available metagenomics data of 278 pigs in the different farms from three countries (Xiao et al., 2016). The antibiotic usage patterns in these pigs have been well-characterized. The pigs in French farms were fed organically and pigs in Danish farms received antibiotics only at weaning, while pigs in the Chinese farm were continuously fed on antibiotics. Metals and biocides were used in all farms. We hypothesized that the use of metals and biocides would exert strong selective pressures on ARGs in intensive farming regardless of antibiotic use.

## Methods

### Sample information, data collection and pre-processing

Table S1 illustrates the antibiotic exposure on 278 pigs in different farms (Xiao et al., 2016). Seventy-eight pigs in the Chinese farm were continuously exposed to various antibiotics, while one hundred pigs in Danish farms were only exposed at weaning. One hundred pigs in French farms were raised completely organic, although a subset of them was raised on farms with previous use of antibiotics.

The clean Illumina Hiseq paired reads of 278 pig gut metagenomes, which have removed adaptor contamination, low-quality reads and pig genomic DNA, were retrieved at https://www.ebi.ac.uk/ena/data/view/PRJEB11755 from European Bioinformatics Institute (EBI). Assembly of contigs was done with megahit (version 1.1.3) with default options (D. Li et al., 2015). Any contigs shorter than 500bp were filtered out. The prediction of open reading frame (ORF) in the contigs was performed with prodigal (version 2.6.2) in META mode (Hyatt et al., 2010).

### ARG, BRG, MRG, and MGE prediction

The antibacterial biocide and metal resistance genes database (BacMet, version 2.0) was used for the predictions of BRG and MRG (Pal et al., 2014): the amino acid sequences of predicted ORFs were subjected to similarity searches against the BacMet database with diamond search in the more sensitive mode and k1 option (Buchfink et al., 2014). Only BRGs and MRGs with at least 90% identity and a maximum e-value of 1 × 10^−3^ were retained. The comprehensive Antibiotic Resistance Database (CARD database, version 3.0.0) was used to detect ARGs: the amino acid sequences of predicted ORFs were aligned with the CARD database through RGI identifier with diamond as aligner (Jia et al., 2017). Only ARGs with “Strict” or “Perfect” significance cut-off were preserved for further analysis. MGE homologs were characterized using PFAM (Finn et al., 2016) and TnpPred (Riadi et al., 2012) databases through HMMER v3.1b2, with an e-value of 1 × 10^−5^ as a threshold (Sáenz et al., 2019). If the predicted ORF had more than one hit for MGE homologs, the hit with the lowest e-value was retained. In the present study, MGEs were classified into 9 categories according to functions: conjugative transposon, integrase/integrase-related, phage integrase, recombinase, resolvase, RteC (related to tetracycline conjugative transposon), transposase/transposase-related, transposition related, transposon breakage related.

### Gene coverage/abundance calculation

Clean reads were aligned against the reference ORF nucleotides with BWA aligner (Li, 2013). The alignment summary statistics were calculated using Samtools idxstats (Li et al., 2009). For the comparisons of gene coverage/abundance between samples, GCPM (Gene Coverage Per Million) value was used to normalize the mapped reads for sequencing depth and gene length. The formula is GCPM 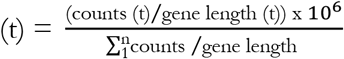 where GCPM (t) is the GCPM value of gene t, counts (t) is the number of mapped reads to the gene (t), gene length (t) is the length of gene (t), n is the number of all the predicted ORFs.

### Statistical analysis and R application

All statistical analysis and data sorting in the study were done in R (R Development Core Team, 2011). Between-individual diversity (β-diversity) of gene coverage/abundance was evaluated by Bray-Curtis dissimilarity matrices through R function “vegdist” in the package “vegan” (Oksanen et al., 2008). β-diversity matrices were ordinated by PCoA plot (R function “plot_ordination” in R package “phyloseq”) (McMurdie and Holmes, 2013). Permutation multivariate analysis of variance (PERMANOVA) was used to compare the differences in β-diversity between groups when FDR correction was required (R function “pairwise.adonis” in R) (Arbizu, 2017). Within-individual diversity (α-diversity) was measured by observed richness of gene coverage/abundance. One-way ANOVA (R package “stats”) was used to compare the differences in α-diversity among groups. TukeyHSD test (R package “stats”) was used to do pairwise comparisons of α-diversity. The pairwise comparison of gene abundance between three groups was done using the Wilcoxon rank sum test (R function “pairwise.wilcox.test”). Venn diagram was plotted using R function “draw.triple.venn” in R package “VennDiagram” (Chen and Boutros, 2011). Fisher’s exact test was used to test the statistical significance of the number of contigs between groups (R function “fisher.test” in R package “stats”).

### Co-occurrence analysis

The resistance genes located in the same assembled contig were considered as co-occurring. Co-occurrence was considered trustable if the same association was present in at least two different contigs.

### Contig source screening

Fifteen million contigs from 138,683 bacterial genomes in the NCBI Reference Sequence Database (RefSeq) were downloaded (19.12.2018) and the downloaded bacterial genomes were indexed to generate a BLAST database with makeblastdb. Blastn (version 2.6.0) was used to search for the contigs carrying ARGs, MGEs, BRGs/MRGs detected in the study from the reference genomes. A minimal blast similarity of 95%, a shortest alignment length of 10kb and a biggest e-value of 10e-10 were used to filter blast hits. R function “heat_tree” in “metacoder” package (Foster et al., 2017) was used to plot the taxonomic tree to visualize the bacterial contig source. The tree branch represents the affiliation relation between taxa.

## Results

### Gut microbiome of pigs in the Chinese farm contained the highest abundance of resistance genes

The total loads of BRGs, MRGs, and potentially mobile ARGs (ARGs found together with MGEs in one contig) varied significantly between different farms (Fig. 1A). When looking at all resistance determinants, the gut microbiome of pigs in the Chinese farm carried the highest load (Pairwise Wilcoxon rank-sum test, FDR-adjusted; Pigs in Danish, French and Chinese farms were represented by D, F and C, respectively. Mobile ARG: C vs D, *P* < 2e-16; C vs F, *P* < 2e-16; D vs F, *P* < 2e-16. BRG: C vs D, *P* = 4.2e-14; C vs F, *P* = 4.5e-10; D vs F, *P* = 0.53. MRG: C vs D, *P* = 5.2e-14; C vs F, *P* = 3.0e-07; D vs F, *P* = 0.027. MGE: C vs D, *P* < 2e-16; C vs F, *P* = 0.073; D vs F, *P* < 2e-16). The pigs in Danish farms had the second-highest abundance of mobile ARGs and similar abundance of BRG to the pigs in French farms. The pigs in French farms had the second-highest abundance of MRGs. Notably, only 10 of 100 pigs in French farms had mobile ARGs in their gut microbiome, though the total abundance of MGEs in French pigs was similar to Chinese pigs.

**Fig. 1.**
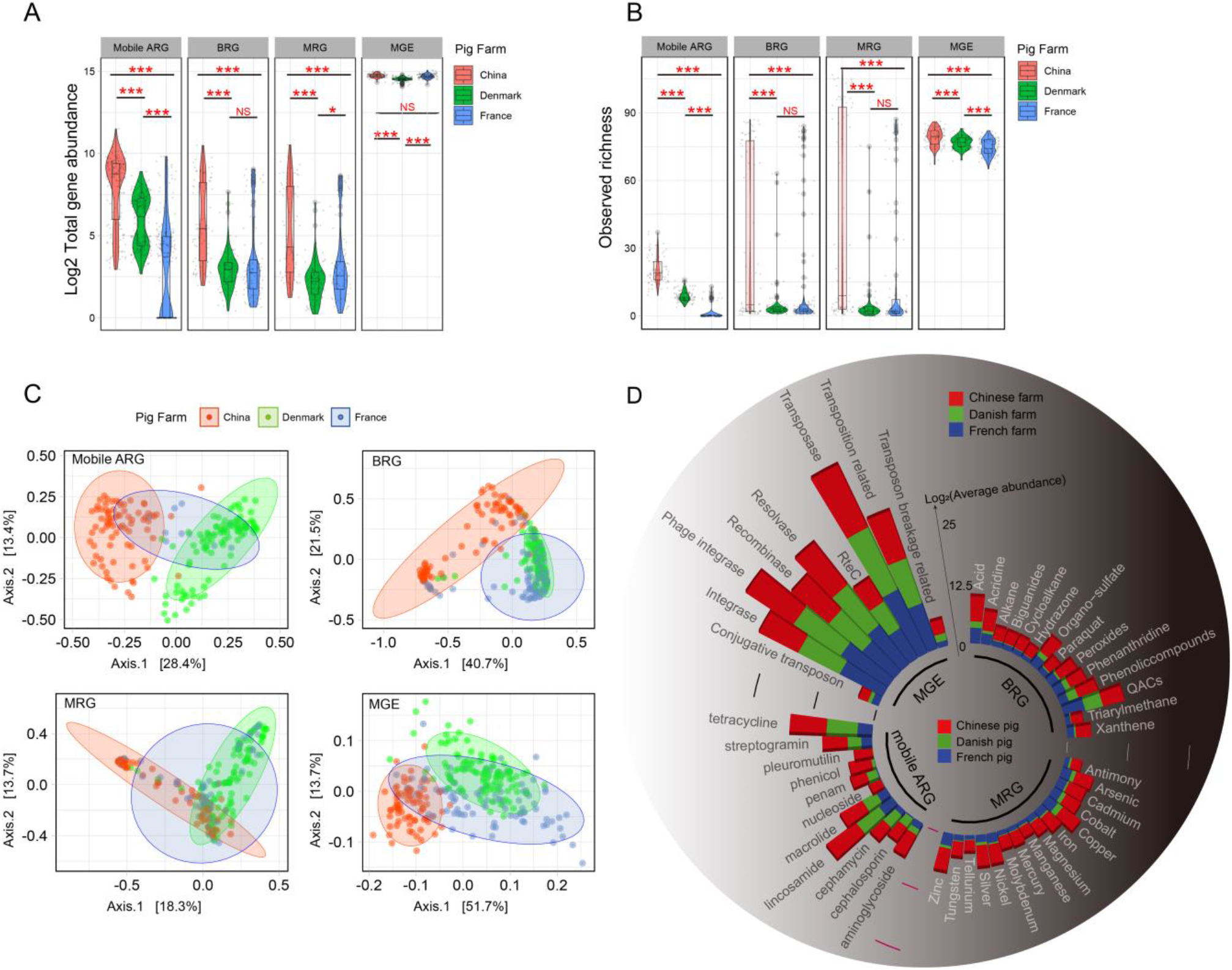
Overview of MRGs, BRGs, mobile ARGs, and MGEs in pig gut microbiomes from different farms. **A).** Violin plot showing the total abundance of resistance genes and MGE per sample. White lines represent the median. Asterisks stand for significant statistical difference between groups (Pairwise Wilcoxon rank-sum test, FDR-adjusted; *P* <0.05*, *P* <0.01**, *P* <0.001***, NS: *P* > 0.05. **B).** Violin plot showing the observed richness of resistance genes and MGE. White lines represent the median. Asterisks stand for significant statistical difference between groups (TukeyHSD test; *P* <0.05*, *P* <0.01**, *P* <0.001***, NS: *P* > 0.05). **C).** PCoA plots showing Bray-Curtis distances of mobile ARG, BRG, MRG and MGE among pigs from Chinese, Danish and French farms. **D).** Average abundance of resistance genes and MGE in the different countries. The abundance of multiple resistance gene was separately counted in the individual function group; for example, the gene with AMR and biocide resistance (BR) was included both in AMR and BR abundance plots. Labels on the stacked bars describe the metals and biocides associated with BRGs and MRGs, different drug classes associated with mobile ARGs and 9 categories of MGEs. Heights of the stacked bars represent the mean value of the total abundance in log-scale.

We calculated observed richness for each resistance determinant in different farms (Fig. 1B). For each gene type, pigs in the Chinese farm were found to have a higher diversity followed by the pigs in Danish farms (TukeyHSD test; Mobile ARG: C vs D, *P* < 1e-07; C vs F, *P* < 1e-07; D vs F, *P* < 1e-07. BRG: C vs D, *P* < 1e-07; C vs F, *P* < 1e-07; D vs F, *P* = 0.11. MRG: C vs D, *P* < 1e-07; C vs F, *P* < 1e-07; D vs F, *P* = 0.08. MGEs: C vs D, *P* = 2.34e-05; C vs F, *P* < 1e-07; D vs F, *P* = 8.28e-05). However, pigs in French farms had similar BRG and MRG diversity compared to pigs in Danish farms.

In general, mobile ARGs, BRGs, MRGs and MGEs had a pronounced separation in composition based on farms (Fig. 1C) (pairwise Adonis test, FDR-adjusted; Mobile ARG/BRG/MRG/MGE: C vs D, *P* = 0.003; C vs F, *P* = 0.003; D vs F, *P* = 0.003). All detected ORFs conferred resistance to 20 metals, 36 biocides, 34 antibiotics, and encoded 9 types of mobile genetic elements. Fig. 1D shows the most abundant genes conferring resistance to 13 metals, 14 biocides, and 10 antibiotics. Overall, of the MRGs detected, genes conferring Zn and Cu resistance were the most abundant. Among BRGs, genes encoding resistance to QACs were the most abundant. When we investigated the potentially mobile fraction of the ARG pool, lincosamide and tetracycline resistance genes were found to be the most abundant and the transposases to be the most common trait related to mobility mode.

### Surveying assembled contigs carrying co-occurrence of ARGs and BRGs/MRGs/MGEs

We then investigated the co-occurrence of resistance genes on contigs, in the categories of ARGs, BRGs/MRGs, and MGEs to evaluate the potential for co-selection (Fig. 2A). Evidently, the pigs in the Chinese farm had the largest amount of overlap from different categories, with 61 contigs harboring all 3 different categories of genes, while this was not observed in any of the pigs in French and Danish farms. MGEs were the largest category and had ample overlap with other categories. The pigs in the Chinese farm had the highest proportion of contigs carrying multiple co-occurrences (Fig. 2B) (Fisher’s exact test; ARGs+MGEs: C vs D, *P* < 2.2e-16, odds ratio = 2.69; C vs F, *P* < 2.2e-16, odds ratio = 18.5; D vs F: *P* < 2.2e-16, odds ratio = 6.87; ARGs+BRGs/MRGs: C vs D, *P* < 2.2e-16, odds ratio = 11.8; C vs F, *P* < 2.2e-16, odds ratio = 2.04; D vs F: *P* < 2.2e-16, odds ratio = 0.17; ARGs+BRGs/MRGs+MGEs: C vs D, *P* < 2.2e-16, odds ratio = NA; C vs F, *P* < 2.2e-16, odds ratio = NA; D vs F: *P* = 1, odds ratio = 0; BRGs/MRGs+MGEs: C vs D, *P* < 2.2e-16, odds ratio = 9.48; C vs F, *P* < 2.2e-16, odds ratio = 36.6; D vs F: *P* = 0.0007, odds ratio = 3.86). The pigs in French farms had a significantly higher proportion of contigs with ARGs and BRGs/MRGs than the pigs in Danish farms. In contrast, the pigs in Danish farms had a higher proportion of contigs carrying resistance genes together with MGEs than the pigs in French farms. Compared to BRGs/MRGs, a larger proportion of ARGs generally tended to sit together with MGEs in the same contigs (Fisher’s exact test; *P* < 2.2e-16, odds ratio: 2.9) (Fig. 2C).

**Fig. 2.**
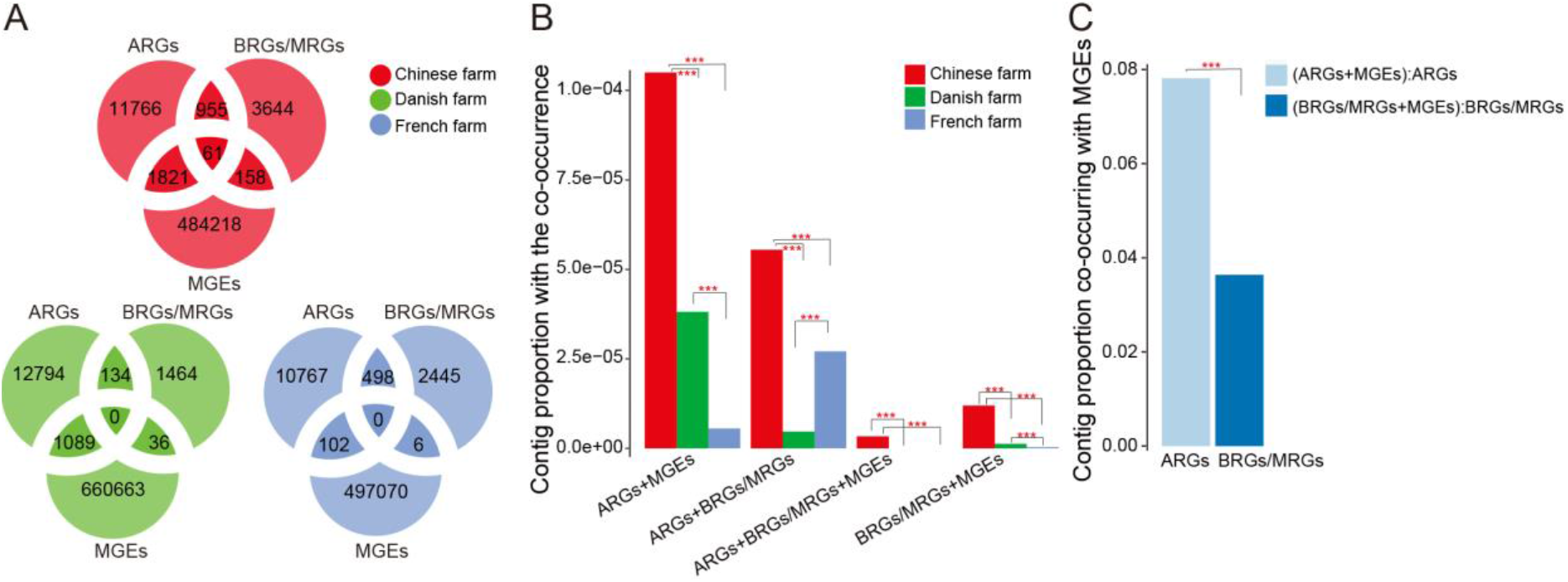
Overview of co-occurrences of ARGs, BRGs, MRGs and MGEs in assembled contigs. **A).** Venn diagram showing the number of contigs carrying ARGs, BRGs/MRGs, MGEs and their combinations in pigs. **B).** The proportion of contigs with co-occurrences of ARGs and BRGs/MRGs, ARGs and MGEs, BRGs/MRGs and MGEs, BRGs/MRGs, ARGs and MGEs in pigs. Asterisks stand for significant statistical difference between groups (Fisher’s exact test; *P* <0.05*, *P* <0.01**, *P* <0.001***). **C).** The proportion of contigs carrying ARGs and BRGs/MRGs with MGEs. Asterisks stand for significant statistical difference between groups (Fisher’s exact test; *P* <0.05*, *P* <0.01**, *P* <0.001***).

### Co-localization of ARGs and BRGs/MRGs/MGEs in assembled contigs

To further verify the co-occurrence of ARGs and BRGs/MRGs/MGEs on the contigs, we visualized the gene organization on representative contigs (Fig. 3A). According to resistances carried in each representative contig, there were six major co-occurrence combinations and 27 co-occurrence subtypes. We selected the contig carrying the most complete gene distribution in each of 27 subgroups as the representative contig. All the co-occurrences involved 131 resistance genes to 17 metals, 28 biocides and 25 antibiotics. Of these, Cu, Ni and Zn resistance genes, fluoroquinolone, penam and tetracycline resistance genes, acid and QAC resistance genes were the most co-occurring genes. The information about these genes has been summarized in Table S2.

**Fig. 3.**
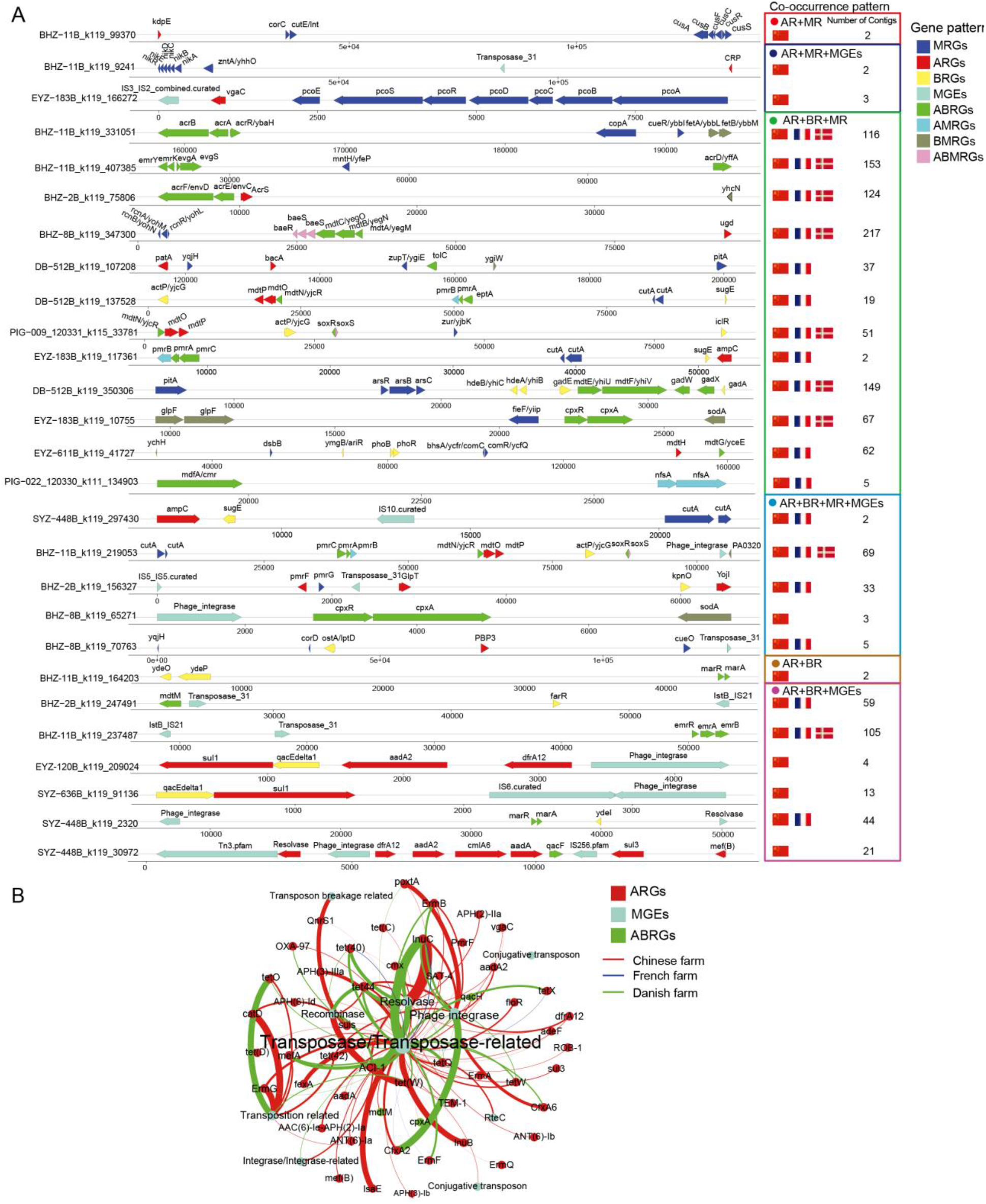
The patterns of Co-occurrence in assembled contigs. **A)** The arrangement of resistance genes within the co-occurrences-carrying representative contigs. Only those co-occurrences present in at least two different contigs are listed. ABRG stands for the gene conveying resistance to antibiotic and biocide; AMRG stands for the gene conveying resistance to antibiotic and metal; BMRG stands for the gene conveying resistance to biocide and metal; ABMRG stands for the gene conveying resistance to antibiotic, biocide and metal. The location and size of genes are proportional to the actual condition; the country flag stands for the location of the farm raising the pigs that have the corresponding co-occurrences and its row order stands for the decreasing co-occurrence frequency. **B)** Network of co-occurring ARGs and MGEs on the same contigs. For view purposes, only the co-occurrences between ARGs and MGEs which are present in at least 5 different contigs are shown in this network. The size of node, node label and the weight of edge are proportional to the number of co-occurred contigs.

Co-occurrences in the pigs from the Chinese farm were detected to be most common, in terms of both the number of contigs carrying co-occurrences and the type of co-occurrence (Fig. 3A). We identified all 27 co-occurrence subtypes in the Chinese farm, 20 subtypes in French farms, and 9 subtypes in Danish farms. Some co-occurrences affiliated with the same subtype were only partly present in the presentative contig. Therefore, to further investigate the abundance of co-occurrences, we plotted a network to show the frequency of co-occurrences between resistance genes in all the contigs (Fig. S1). The detailed information for the network has been summarized in Table S3. As shown in Fig. S1, ARGs in the pigs from Danish farms mostly co-occurred with BRGs, which mostly tended to be functionally associated, such as multidrug efflux transporter genes *acrAB* and its regulator gene *acrR*. Notably, we detected an assembled contig carrying one colistin resistance gene *MCR*-4 and Cd/TBT resistance gene *ygiW* in the pigs from Danish farms.

Some co-occurrences were only detected in the pigs from the Chinese farm (Fig. 3A) – for example, the co-occurrence of fluoroquinolone ARG *CRP* with the Ni resistance operon *nikA/B/C/D/E/R* and Zn resistance gene *zntA*; the co-occurrence of aminoglycoside ARG *kdpE* with nine Cu resistance genes *cusA/B/C/F/R/S*, *cutE/F*, *corC*; and the co-occurrence of streptogramin ARG *vgaC* with seven Cu resistance genes *pcoA/B/C/D/E/R/S*. Two archetypal Class I clinical integrons carrying 3’ conserved segment (CS) of an antiseptic-resistance gene *qac*EΔ1, a sulfonamide resistance gene *sul*1, 5’ CS of the integrase, as well as gene cassettes *aad*A2-*dfr*A17 (Integron A) or an IS6 transposase gene (Integron B) were only detected in the pigs from the Chinese farm. An analogous structure consisted of *qacF*, *sul*3, an integrase, and gene cassette *dfr*A12-*aad*A2-*cml*A6-*aad*A was only detected in the pigs from the Chinese farm as well (Integron C). To verify whether these three integrons are located on plasmids, we mapped the three representative contigs against the PLSDB plasmid database (Galata et al., 2019). It turned out that Integron C and Integron B were very similar to DNA fragments in 180 and 73 plasmids, respectively (Mash search, Max distance=0.1). Integron C was frequently found in *E. coli*, *Klebsiella pneumonia* and *Salmonella enterica* (Fig. S2-a and Table S4). The bacteria harboring Integron B covered a wide range of species such as *E. coli*, *Pseudomonas aeruginosa, Klebsiella pneumonia* and *Enterobacter cloacae* (Fig. S2-c and Table S5). The gut microbiome in the pigs from the Chinese farm harbored some co-occurrences related to polymyxin including colistin resistance genes. For example, the polymyxin ARG *ugd* was located in the same contig with Co-Ni-Fe related resistance genes *rcnA/B/R*, multidrug efflux system MdtA/B/C, BaeS/R two-component system; Polymyxin resistance genes *pmrA/B/C*, colistin heteroresistance related *soxR/S*, co-occurred with Cu resistance gene *cutA*, acid resistance gene *actP*, Cd-Hg-TBT-H_2_O_2_-HCl resistance gene *PA*0320 and a phage integrase.

Some co-occurrences were detected in pigs from both French and Chinese farms. For example, the polymyxin resistance gene *pmrF*, fosfomycin resistance gene *GlpT*, microcin J25 resistance gene *YojI* were found on the same contig with a QACs- H_2_O_2_ resistance gene *kpnO* and a Fe resistance gene *pmrG*; The genes encoding the two-component regulatory system CpxAR co-occurred with Fe-Zn resistance gene *fieF*, Se-H_2_O_2_ resistance gene *sodA* and Sb-As-glycerol uptake and resistance gene *glpF*; Cu resistance gene *copA*, *cueR* and Fe-H_2_O_2_ resistance genes *fetA*, *fetB* were frequently detected to co-occur with genes encoding the AcrR/A/B multidrug efflux pump operon; And acid resistance genes including *gadE*, *hdeA*, *hdeB*, *mdtE*, *mdtF*, *gadW*, *gadX*, *gadA* were located in the same contig with Zn-Te resistance gene *pitA* and As resistance gene *arsA/B/R*; Beta-lactam resistance gene *PBP*3 was found to co-occur with two Cu resistance gene *corD*, *cueO*, Fe resistance gene *ygjH* (only in Chinese pigs), one BRG *ostA* and one transposase gene; A Cd-H_2_O_2_-HCl resistance gene *yhcN* co-occurred with the efflux pump system AcrE/F and its repressor AcrS; A broad spectrum MDR ABRG *mdfA* was detected in the same contig with Cr-nitrofuran resistance gene *nfsA*.

In this study, we plotted a network to demonstrate the co-occurrences between ARGs and MGEs in the same contigs (Fig. 3B). In total, 282/1261 ARGs were co-located with MGEs on the same contigs, referred to as mobile ARGs. Mobile ARGs are abundant in the pigs from Chinese and Danish farms, especially transposase. The mobile lincosamide resistance gene *lnuC* was most detected in all three farms. The mobile cephamycin resistance genes *CfxA*2*/CfxA*6 and penam resistance gene *ACI*-1 were also abundant in the pigs from Danish and Chinese farms. Some tetracycline resistance genes (*tet*(40)/(42)/*(C)/(D)*/44/*W/O/Q/X/adeF*), macrolide resistance genes (*ermA/B/F/G/Q*) and aminoglycoside resistance genes (*ANT*(6)-*Ia/ANT*(6)-*Ib/APH*(2)-*IIa/APH*(3)-*Ib/APH*(3)-*IIIa/APH*(6)-*Id/aadA/aadA*2) were also frequently detected to co-locate with MGEs in the same contigs. The polymyxin resistance gene *pmrF* was found to be in the vicinity of genes encoding transposases in the same contig. Colistin resistance-related genes *mexB* and *oprM*, and polymyxin B resistance gene *eptA* were also detected to co-occur with MGEs in Chinese and Danish pigs. The detailed information for the network has been summarized in Table S6.

### Mobilizable contigs carrying ARG, BRG/MRG and MGEs are concentrated in *Enterobacteriaceae*

To investigate which clades of bacteria tended to maintain and spread ARGs in the form of co-selection and also confirm the accuracy of contig assembly, we blasted the contigs carrying ARGs, MGEs, BRGs/MRGs against the NCBI RefSeq database consisting of 138, 683 genomes. All the contigs could be found in reference genomes. The tree in Fig. 4 demonstrated that those mapped bacteria and the size of nodes was proportional to the mapped occurrence of the bacterial taxa. Proteobacteria was the most often hit phylum, under which family *Enterobacteriaceae* had the biggest class branch. Notably, the opportunistic pathogen *E. coli* and the pathogen *Shigella flexneri* were the most mapped species (Fig. 4).

**Fig. 4.**
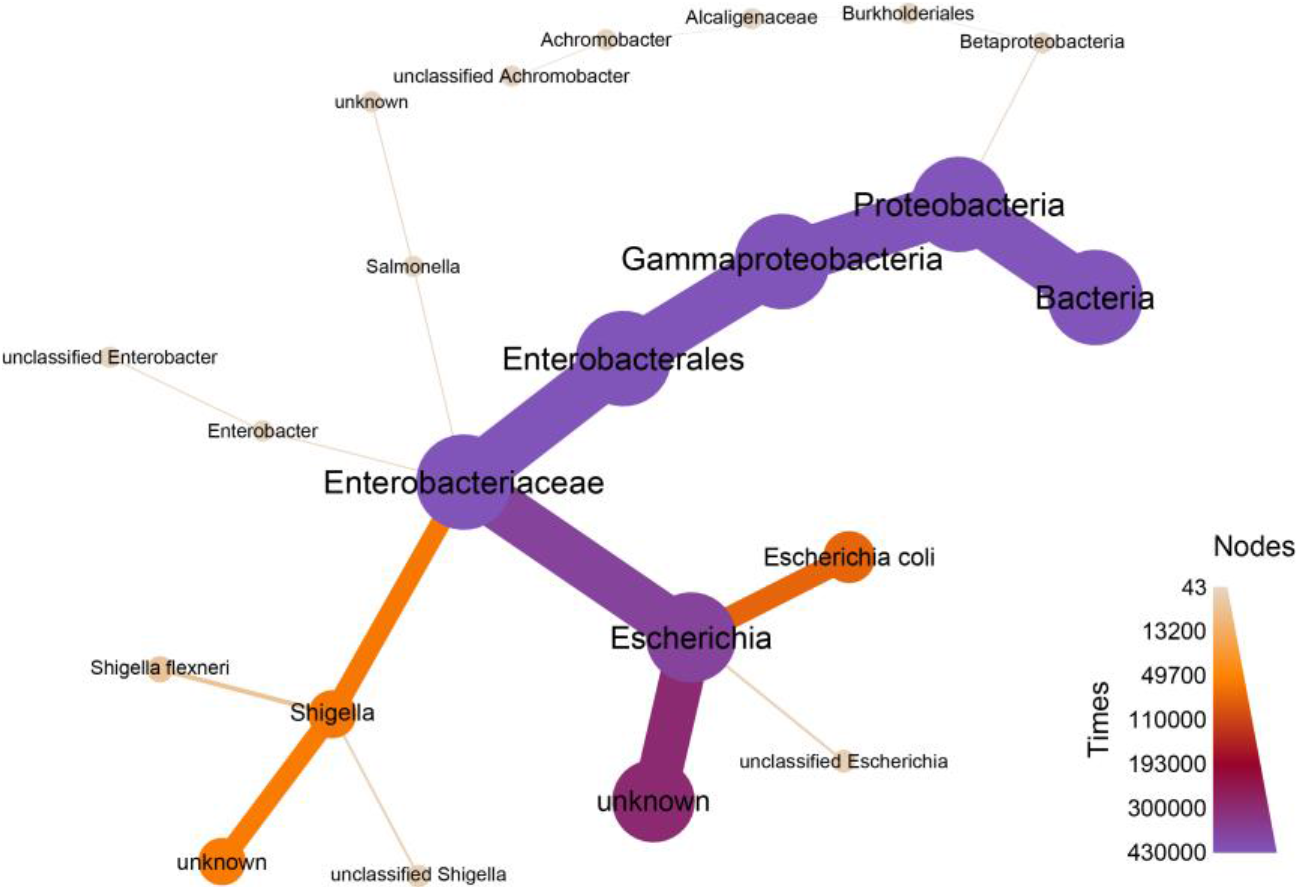
Tree showing bacteria harboring contigs with ARGs, MGEs, MRGs/BRGs. The node size, node label size, and edge weight were proportional to mapped frequency. All the bacteria taxa that had a successful match have been shown in Table S7.

## Discussion

The current study provides a comprehensive analysis based on the genetic linkage of bacterial co-occurrence profiles between ARGs and other genetic elements including MRGs, BRGs and MGEs in the gut microbiome of 278 pigs from Danish, French and Chinese farms. The antibiotic feeding mode of pigs in the Chinese, Danish and French farms can stand for an unmonitored mode, European standard mode and organic mode, respectively. Compared to antibiotics, the usage of metals as a growth promotor in pig farming has received little restriction around the world. But even worse, unmonitored biocide usage as a disinfectant in the pigsty and its residual in vegetable feed did so far not cause much attention. In this study, we found a positive association between the extent of antibiotic use and the load of mobile ARGs. The gut of pigs in the Chinese farm contained the most abundant mobile ARGs, followed by pigs from Danish farms and lastly the pigs from French farms. The abundance and type of MRGs and BRGs did not vary a lot among the different farms. Pigs in all farms had abundant MRGs and BRGs, especially genes encoding resistances for Zn, Cu, Ag, Ni, As, Cd, QAC. In general, Cu and Zn are the most common growth promoters in animal feeding and residuals can be always found in manure (Mazhar et al., 2021). QAC resistance BRG was frequently found in the pigs from all farms. Over the past decade, the use of QACs as detergent, disinfectant and preservative has dramatically increased in industry, hospitals, and cosmetics (Buffet-Bataillon et al., 2012), accompanied by an elevated occurrence of QAC resistance genes in many bacteria (Bischoff et al., 2011; Heir et al., 1999; Langsrud et al., 2003; Seier-Petersen et al., 2015; Zmantar et al., 2011). Unfortunately, we cannot provide an effects size estimate using our study, since no quantitative or qualitative data on the use of these compounds was collected. In addition to the widespread use of QACs, the spread of these resistance genes via HGT among prokaryotes may be another reason for their widespread presence since a significant correlation between QAC resistance genes and MGEs was detected in this study. The spread of resistance determinants through HGT is a favored way for microbes to adapt to complex environmental pressures (Ye et al., 2017). In this study, around 10.8% (282/1261) ARGs, which co-occurred with MGEs especially genes encoding transposase, had the potential to be transferred via HGT. Some previous work had shown that the levels of transposases were highly correlated to the abundance of ARGs from other environments (Aziz et al., 2010; Zhu et al., 2013).

In this study, co-selection phenomena were investigated via co-occurrence patterns as a proxy. We found that regardless of antibiotics use or little/no antibiotic use, AMR seems to be maintained by co-occurrence with MRGs/BRGs. Despite little/no antibiotic use in Danish and French farms, cross-resistance, for example, *mdtA/B*, *mdtE/F*, *mdtN/O*, *cpxA/R*, *acrA/B/R*, *acrE/F*, *emrA/B/R*, *emrK/Y*, *gadA/X/W*, *baeR/S*, *soxS/R*, which have multidrug resistance towards antibiotics, biocides and/or metals, can still help to maintain AMR under exposure stress from biocides and/or metals. Of these, some multidrug resistance genes can encode multiple-component regulatory systems that mediate co-regulation as well. For example, the BaeRS two-component regulatory system can activate transcription of many resistance genes (Nishino et al., 2005) such as *mdtA/B/C/D* transporter gene clusters and *acrD* (Baranova and Nikaido, 2002; Nagakubo et al., 2002). The CpxAR two-component regulatory system can activate transcription of the multiple antibiotic resistance regulatory operon *marA/B/R* (Weatherspoon-Griffin et al., 2014). *soxS* can enhance the expression of the Zn uptake system ZnuACB (Warner and Levy, 2012) and the AcrAB-TolC multidrug efflux protein complex (Pérez et al., 2012). In addition to cross-resistance, co-resistance phenomena detected in the pigs especially from French farms further highlighted the role of metals/biocides on the maintenance and spread of ARGs. These co-resistance patterns were either functionally connected such as in *acrA/B/R* or probably reflected the historical and current chemical environments (Ye et al., 2017). Zhu *et al* found that the co-selection of Cu, Zn, tetracycline resistance determinants and MGEs would be favored in exposed microbial communities due to the use of Cu, Zn, and antibiotics such as tetracycline in pig farming (Zhu et al., 2013). Accordingly, we speculate that microbes in the gut of pigs faced the selection pressures from metals, antibiotics and biocides including Cu, Ni, Zn, fluoroquinolone, penam, tetracycline, cephalosporin, Acid, QACs, Acridine, *etc*. In summary, antibiotics are not the sole factor for the spread and preservation of ARGs in their ecological environment.

Notably, we found some polymyxin resistance genes and their co-occurrences with other resistance genes. As we know, polymyxins have reemerged as a final line of defense against Gram-negative ‘superbugs’ (Sun et al., 2018). The co-occurrences of polymyxin resistance genes and other resistance genes would aggravate their spread and maintenance. In this study, although we found integrons carrying ARG cassette in pigs only from the Chinese farm, the plasmids carrying these integrons have been found around the world (Fig. S2-b/d) and mostly in pathogens. The co-occurrence between QAC resistance genes *qacF*/*qacE*Δ1 and ARG cassettes in the integrons could help AMR maintenance in pathogens. Similarly, Gaze *et al* found that the incidence of Class I integrons was significantly higher for bacteria exposure to QACs (Gaze et al., 2005).

Enterobacteriaceae tend to have more ARGs/MRGs/MGEs-carrying DNA fragments, especially in *E. coli* and *Shigella flexneri*. *E. coli* is known to carry a high degree of ARGs (Li et al., 2021) and *Shigella flexneri* can cause a variety of communicable bacterial dysenteries in their hosts (Jennison and Verma, 2004). The era of plentiful antibiotics is forcing more bacteria to develop ARGs to survive and grow in the newly established toxic environment, while non-degradable metals and easily accessible biocides probably further promote the maintenance and spread of AMR in the form of co-occurrence investigated in this study, which represents an increased public health risk.

While this study can distinguish patterns of co-selection that align with countries and farms that are known to have large differences in antibiotics usage, it is a limitation that there are no observations on many other aspects of farming practices. Cleaning agents and frequency would be particularly pertinent given the observed co-selection of biocide resistance genes, but other differences in cultural and regulatory practices could impact both the microbiome and the resistome this can harbor. Ideally, observations of fully factorial designs could be used to separate the variance of these, but ultimately intervention designs are needed to establish practices that can minimize the horizontal spread of ARGs.

## Conclusion

In this study, we have used publicly available data to map the mobilizable resistance genes in pig gut microbiomes and exploited the patterns of co-selection by non-antibiotic factors. The genetic evidence presented here clearly suggests these factors contribute to the maintenance of AMR in pig farming. We demonstrate the commonness of this co-selection with genetic evidence and augment this with an overview of mobilizing elements. We clarify that this maintenance is not a random selection from a mobilized pool but pertains to specific phylogenetic clades. We hope this work will give further insights into the genetics of co-selection and the implications of non-pharmaceutical antibiotic agent usage in the propagation of AMR. Specifically, this work illustrates the need for a comprehensive survey of co-selection potential for effective agronomic practices policymaking aiming for a reduction of the global AMR burden in a One Health approach.

## Acknowledgments

We thank Chinese scholarship council and Novo Nordisk Foundation for funding support. ADB was supported by the Danish Council of Independent Research (grant 6111-00471B) and Lundbeck Foundation (grant R140–2013-13528).

## CRediT authorship contribution statement

**Xuanji Li:** Data collection, Project administration, Analysis, Writing - original draft. **Christopher Rensing:** Project administration, Writing - review & editing. **Gisle Vestergaard:** Methodology, Writing - review & editing. **Joseph Nesme:** Project administration, Writing - review & editing. **Shashank Gupta:** Project administration, Writing - review & editing. **Manimozhiyan Arumugam:** Methodology, Writing review & editing. **Asker Daniel Brejnrod:** Data collection, Supervision, Analysis, Writing - review & editing. **Søren Johannes Sørensen:** Supervision, Writing - review & editing, Funding acquisition.

