## Supplementary material for "Metagenomic evidence for co-occurrence of antibiotic, biocide and metal resistance genes in pigs": Fig.S1

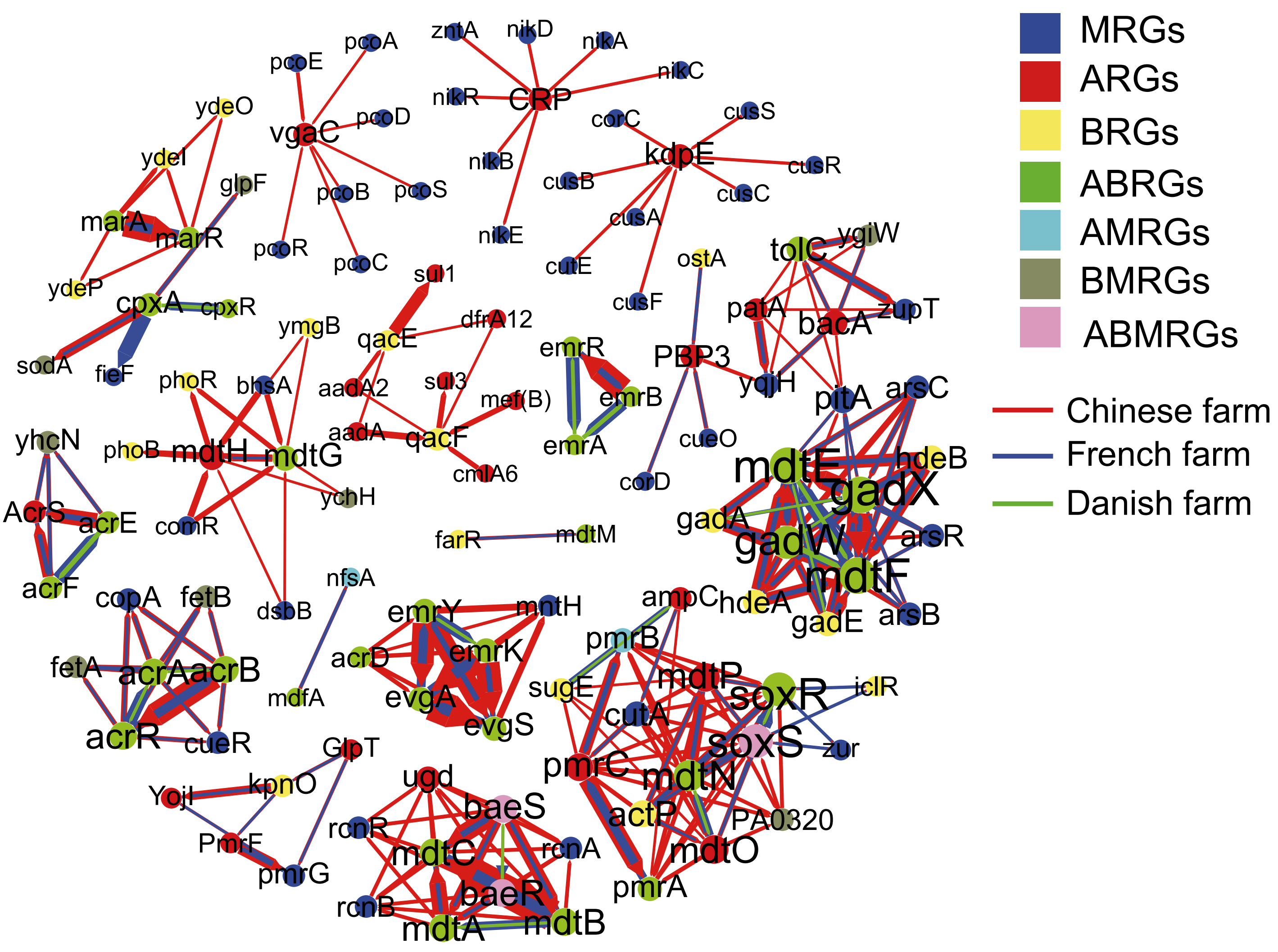


**Fig. S1.** Network of co-occurrence between ARGs and BRGs/MRGs. This network summarized and quantified the co-occurrences between ARGs and BRGs/MRGs in the three farms.
