## Supplementary material for "Metagenomic evidence for co-occurrence of antibiotic, biocide and metal resistance genes in pigs": Fig.S2

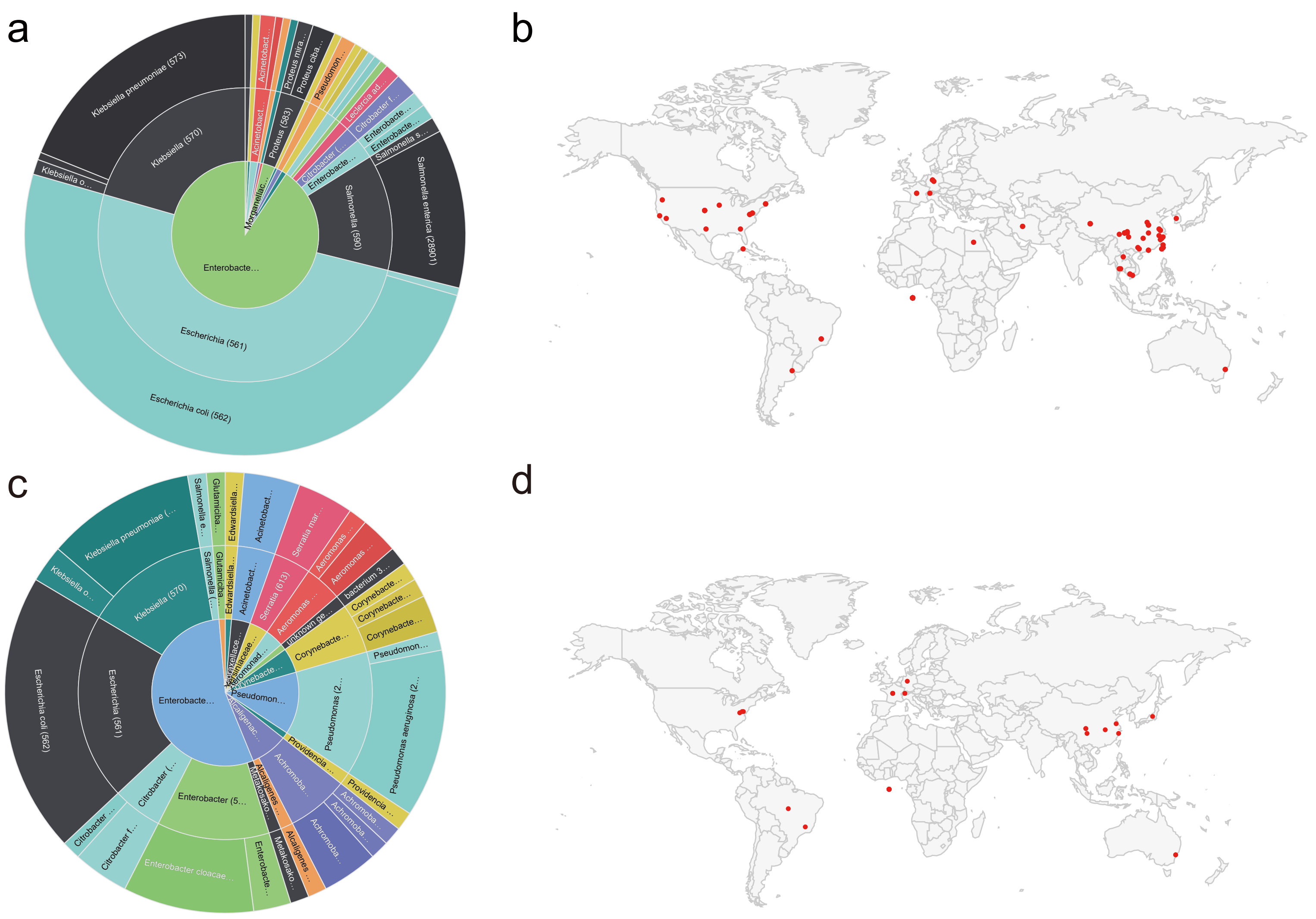


**Fig. S2**. Bacterial taxonomy and sampling location of mapped plasmids with Integron C (a,b) and Integron B (c,d) against PLSDB database.
